# Comparative assessment of the effects of bumped kinase inhibitors on early zebrafish embryo development and pregnancy in mice

**DOI:** 10.1101/2020.06.17.157529

**Authors:** Nicoleta Anghel, Pablo A. Winzer, Dennis Imhof, Joachim Müller, Javier Langa, Jessica Rieder, Lynn K. Barrett, Rama Subba Rao Vidadala, Wenlin Huang, Ryan Choi, Mathew A. Hulverson, Grant R. Whitman, Samuel L. Arnold, Wesley C. Van Voorhis, Kayode K. Ojo, Dustin J. Maly, Erkang Fan, Andrew Hemphill

## Abstract

Bumped kinase inhibitors (BKIs) are effective against a variety of apicomplexan parasites. Fifteen BKIs with promising *in vitro* efficacy against *Neospora caninum* tachyzoites, low cytotoxicity in mammalian cells, and no toxic effects in non-pregnant BALB/c mice, were assessed in pregnant mice. Drugs were emulsified in corn oil and applied by gavage for 5 days. Five BKIs did not affect pregnancy, 5 BKIs exhibited 15-35% of neonatal mortality, and 5 compounds caused strong effects (infertility, abortion, stillbirth and pup mortality). Additionally, the impact of these compounds on zebrafish (*Danio rerio*) embryo development was assessed by exposing freshly fertilized eggs to 0.2-50μM of BKIs and microscopical monitoring of embryo development in a blinded manner during 4 days. We propose an algorithm that includes quantification of malformations and embryo deaths, and established a scoring system that allows to calculate an impact score (S_i_) that indicates at which concentrations BKIs visibly affect zebrafish embryo development. Comparison of the two models showed that for 9 compounds no clear correlation between S_i_ and pregnancy outcome was visible. However, those 3 BKIs affecting zebrafish embryos only at high concentrations (40μM or higher) did not impair mouse pregnancy at all, and those 3 compounds that inhibited zebrafish embryo development already at 0.2μM showed detrimental effects in the pregnancy model. Thus, the zebrafish embryo development test has a limited predictive value to foresee pregnancy outcome in BKI-treated mice. We conclude, that maternal health-related factors such as cardiovascular, pharmacokinetic and/or bioavailability properties also contribute to BKI-pregnancy effects.

## 1. Introduction

*Neospora caninum* and *Toxoplasma gondii* are obligate intracellular protozoan parasites which belong to the phylum Apicomplexa. They have a worldwide distribution and are found in many different species, infecting different cell types and tissues. *Toxoplasma* and *Neospora* undergo three distinct infective stages: rapidly proliferating tachyzoites that cause acute disease, and are responsible for vertical transmission during pregnancy; slowly proliferating bradyzoites that are responsible for chronic infection, which are enclosed in tissue cysts that are orally infective; and sporulated oocysts containing sporozoites, the end product of a sexual cycle that takes place within canine (*Neospora*) or feline (*Toxoplasma*) intestinal tissue, which are shed with the feces and ingested by intermediate hosts [1,2].

Primary oral infection through oocysts or tissue cysts from both species during pregnancy can result in exogenous transplacental infection of the fetus, with grave consequences. In cattle persistently infected with *N. caninum*, endogenous transplacental infection upon recrudescence and reactivation of dormant bradyzoites during pregnancy represents the most abundant mode of fetal infection. Thus, *N. caninum* causes bovine abortion, and inflicts significant economic losses, up to 1.3 billion US dollars according to data derived from 10 countries [3]. N. caninum infection has also been reported in many other species including sheep and goats, but not in humans. *T. gondii* poses highly important veterinary, medical, economic, and public health problems, in that it infects all warm-blooded animals, but also has a very high zoonotic potential [1]. In humans, congenital toxoplasmosis causes fetal malformations and abortion, and recrudescence of latent infection or new infections in immune-suppressed individuals can cause acute toxoplasmosis, with potentially lethal outcomes [4]. Calcium-dependent protein kinase 1 (CDPK1) has emerged as an important drug target in apicomplexans [5]. Homologues of CDPK1 are not present in mammals, but appear to have descended from the plant kingdom [6]. CDPK1 is crucially involved in signaling events that lead to host cell invasion and egress [7, 8]. A promising group of compounds are bumped kinase inhibitors (BKIs), which target CDPK1. CDPK1 is structurally similar in many apicomplexans [5]. The high degree of efficacy and specificity of BKIs in apicomplexans relative to mammalian kinases is due to differences in the topology of a hydrophobic pocket within the ATP binding site that is, in part, determined by the size of their respective gatekeeper residues [9]. BKIs have proven *in vitro* activity against Babesia bovis [6] and *Besnoitia besnoiti* [10], as well as *in vitro* and *in vivo* against *Cryptosporidium* spp. [11–17], *Cystoisospora suis* [18], *T. gondii* [19–21] and *N. caninum* [21–23]. Despite promising efficacies, some BKIs have shown toxicity *in vivo*. For instance, BKI-1294 inhibited the human Ether-à-go-go Related Gene (hERG) *in vitro* [24]. The AC (5-aminopyrazole-4-carboxamide) compound BKI-1517 did not inhibit hERG *in vitro*, but caused a significant change in mean arterial pressure, when dosed in rats. However, in pregnant BALB/c mice infected with *N. caninum*, BKI-1294 had a profound impact on vertical transmission and offspring mortality [25]. Similarly, BKI-1294 impacted on *T. gondii* oocyst infection in pregnant CD1 mice, achieving 100% offspring survival and significant inhibition of vertical transmission [20]. Two other BKIs, −1517 and −1553, also inhibited *N. caninum* proliferation *in vitro*, and were highly efficacious and safe in non-pregnant mice, but were shown to interfere in pregnancy outcome in *N. caninum* infected mice [23]. Thus, potential pregnancy interference is an important issue for the development of drugs designed to inhibit vertical transmission. This interference can be assessed by performing a small pilot study with few non-infected, pregnant mice, which are treated with the drug during early-to-mid-pregnancy [26].

Zebrafish (*Danio rerio*) have been increasingly used for the evaluation of agrochemical agents and toxicity assessments of pharmaceuticals [27, 28]. Zebrafish embryos are a valuable model for toxicity testing and have numerous advantages: low maintenance cost, small size, transparency of the embryo, and rapid development, all of which render investigations time and cost-effective [27, 29, 30]. The organization for economic cooperation and development (OECD) designed a fish embryo toxicity test using the zebrafish model. This assay is used to determine the lethal effects of chemicals on embryonic stages of fish as an alternative method to acute toxicity tests with juvenile and adult fish [31].

In the present study, a zebrafish embryo development test was established to assess the toxicity of 15 distinct BKIs with respect to early embryonal death and induction of malformations, and to investigate the predictive value of the BKI-zebrafish test in relation to pregnancy interference in mice. We show here that the zebrafish embryo model can be a cost-effective screen of novel drugs that can contribute to the reduction of higher-order animals to be used in such studies.

## 2. Materials and methods

### 2.1 Tissue culture, biochemicals and compounds

Cell culture media was purchased from Gibco/BRL (Zürich, Switzerland), and biochemical reagents were obtained from Sigma (St. Louis, MO, USA). The BKIs (Table 1) were obtained from the Center for Emerging and Re-emerging Infectious Diseases (CERID), University of Washington, Seattle, USA. For *in vitro* studies and use in zebrafish assays, BKIs were stored as stock solutions of 50mM in dimethyl sulfoxide (DMSO) at −20°C. For application in mice, BKIs were directly suspended in corn oil and administrated by oral gavage.

**Table 1.** Structures of BKIs investigated in this study and their corresponding effects on *N.caninum* proliferation and cytotoxicity in HFF, HEPG2 and CRL-8155 cells. ^+^and^++^ indicate that EC_50_ data are from [25] and [23] respectively.

| BKI | Structure | Proliferation EC<sub>50</sub> (μM) | Cytotoxicity CC<sub>50</sub> (μM) |  |  |
| --- | --- | --- | --- | --- | --- |
|  |  | *N. caninum* | HEP-G2 | CRL-8155 | HFF |
| 1294 |  | 0.040<sup>+</sup> | >40 | >40 | >20 |
| 1369 |  | 0.035 | >80 | >80 | >20 |
| 1770 |  | 0.144 | >80 | >80 | >20 |
| 1750 |  | 0.076 | >40 | >40 | >20 |
| 1772 |  | 0.112 | >80 | >80 | >20 |
| 1553 |  | 0.180<sup>++</sup> | >40 | 39.5 | >20 |
| 1757 |  | 0.142 | >80 | >80 | >20 |
| 1660 |  | 0.164 | >40 | >40 | >20 |
| 1713 |  | 0.098 | >40 | >40 | >20 |
| 1774 |  | 0.066 | >80 | nd | >20 |
| 1517 |  | 0.269 | >40 | >40 | >20 |
| 1796 |  | 0.067 | >80 | >80 | >20 |
| 1751 |  | 0.117 | >40 | >40 | >20 |
| 1649 |  | 0.071 | >40 | >40 | >20 |
| 1748 |  | 0.068 | >40 | >40 | >20 |

### 2.2 Cytotoxicity assessment in human foreskin fibroblasts (HFF), and human hepatocyte (HepG2) and lymphocyte(CRL-8155) cell lines

HFF were cultured in Dulbecco's modified Eagle (DMEM) with phenol red, supplemented with 10% fetal calf serum (FCS) and 50U of penicillin/ml and were maintained as previously described [32]. For measuring cytotoxicity, HFF monolayers were grown in 96 well plates and were exposed to medium containing serial dilutions of BKIs (0.0009-20 μM). Controls included cultures treated with DMSO, and HFF in medium alone. After culture at 37°C/5%CO_2_ for 3 days, medium was discarded and cells were washed once with PBS. Cytotoxicity was measured by Alamar blue assay as described [32]. HepG2 (hepatocyte) or CRL-8155 (lymphocyte) cells were cultured as previously described [33] with minor changes. HepG2 cells were grown in DMEM/F12 medium plus L-glutamine and 15mM HEPES, CRL-8155 cells in RPMI-1640 medium plus 10mM HEPES, 1mM sodium-pyruvate, and L-glutamine, both media were supplemented with 10% fetal bovine serum. Cells were grown in the presence of varying concentrations of inhibitor, starting at 40μM or 80μM, depending on the solubility, and then serial 3-fold dilutions were performed. Cultures were maintained for 48h at 37°C and 5%CO_2_ in 96-well flat-bottom plates. Growth was quantified using Alamar Blue (Life Technologies, USA). Percent growth inhibition (CC_50_ = drug concentration that causes 50% cytotoxicity) by test compounds was determined based on cultures incubated with DMSO negative and quinacrine positive controls (0% and 100% growth inhibition, respectively) [33]. All assays were performed in triplicate.

### 2.3 In vitro efficacy against N. caninum

A transgenic *N. caninum* strain constitutively expressing beta-galactosidase (Nc-βgal) [25] was maintained by serial passages in HFF. HFF monolayers cultured in 96-well plates were infected with 10^3^ Nc-βgal tachyzoites/well in the presence of serial dilutions of BKIs (0.0164-2μM). After 3 days of culture in the presence of the drugs, wells were washed once with 200μl PBS, and proliferation was assessed as described earlier [25] EC_50_ values (= the half-maximal drug concentration that inhibits tachyzoite proliferation by 50%) were calculated after the logit-log transformation of relative growth and subsequent regression analysis by use of the corresponding software tool contained in the Excel software package (Microsoft, Seattle, WA).

### 2.4 Ethics statement

All protocols involving animals were approved by the Animal Welfare Committee of the Canton of Bern under the license BE101/17. Animals were handled in strict accordance with practices made to minimize suffering. All mice were maintained in a common room under controlled temperature and a 14h/10h light-cycle. All procedures were carried out according to the guidelines set by the animal welfare legislation of the Swiss Veterinary Office.

### 2.5 Assessment of BKI-interference in pregnancy

98 BALB/c female and 49 male mice, 8 weeks old, were purchased from Charles River, Sulzfeld, Germany. Female mice were randomly allocated to 16 experimental groups (6-7 mice/group; 2-3 females/cage; 3 cages/group), with 15 groups undergoing BKI-treatments, and one group receiving placebo (corn oil). Females were oestrus-synchronized by the Whitten effect [22] and were mated for three nights (1 male per 2 females), after which males were removed from the cages. All BKIs (see Table 1) were suspended in corn oil and administered by oral gavage (50mg/kg/day in 100μl) from day 9 to day 13 post-mating as described earlier for BKI-1294 treatment [20, 22, 23]. The only exception is BKI-1796, which was administered at 25mg/kg body weight on days 1, 3 and 5. The placebo group received 100μL of corn oil only/day for 5 days. Weight measurements of each mouse were carried out every 3-4 days to detect pregnant animals and animals were inspected daily to detect potential abortions, which would also cause rapid weight loss. On day 18 post-mating, pregnant females were separated into individual cages, where they gave birth on days 20-22. Live and stillborn pups were counted, and dead pups were removed. Data on fertility, litter size, pup mortality and the clinical status of the dams were recorded. Dams and live pups were monitored at least 2 weeks after birth to rule out possible late adverse effects. At the end of the experiment, pups were euthanized by decapitation. Adult mice were euthanized in a euthanasia chamber using isoflurane and CO_2_.

### 2.6 Assessment of BKI-interference in zebrafish embryo development

The overall setup of the assay is shown in Fig. 1A. At 3h post fertilization (3hpf), zebrafish eggs were transferred into a petri dish (Sarstedt Inc, Nümbrecht, Germany) containing E3-medium (5mM NaCl, 0.17mM KCl, 0.33mM CaCl_2_, and 0.33mM Mg SO_4_, buffered to pH7.5 using sodium-bicarbonate)/osmosis water) containing the tested drug concentration. From there, viable eggs were transferred into the respective 24-well plate (Sarstedt), one egg/well in 1 ml of the same solution. The plate maps are shown in Fig. 1B. Each 24-well plate contained 20 wells with the test solution at a given concentration, and 4 wells without drug as internal controls (iC). The negative control plate (nC) was comprised of 20 wells with 1mL E3 medium/osmosis water. The solvent control (sC) plate consisted of 20 wells comprised of 1mL E3 medium containing 0.01%DMSO. The positive control (pC) plate consisted of 20 wells containing 1mL of 3,4-dichloroaniline (4g/L), which is an established and widely used control substance in zebrafish embryo toxicity tests [34]. The BKI-plates (T) were composed of 20 wells containing 1mL/well of the respective BKI. All BKIs were assessed at 6-8 concentrations, ranging from 0.2 to 50μM. All plates were covered with sealing foil, and embryos were maintained at 28°C. Test solutions were replaced every 24h with fresh solutions. Embryos were viewed and malformations or viability changes assessed in a blinded manner using a Nikon eclipse TS100 light microscope at 10x magnification at 24, 48, 72 and 96hpf, and plates were kept on a heating pad at 26°C during observation and medium changes. The parameters that were monitored and scored are detailed in Fig. 2. At 96hpf, all embryos were euthanized by immersion into a solution of pre-cooled 3-101 aminobenzoic acid ethyl ester (100 μg/L; MS222, Argent Chemical Laboratories, Redmond, WA, USA) and placing them at −20°C for 24 h.

**Figure 1.**
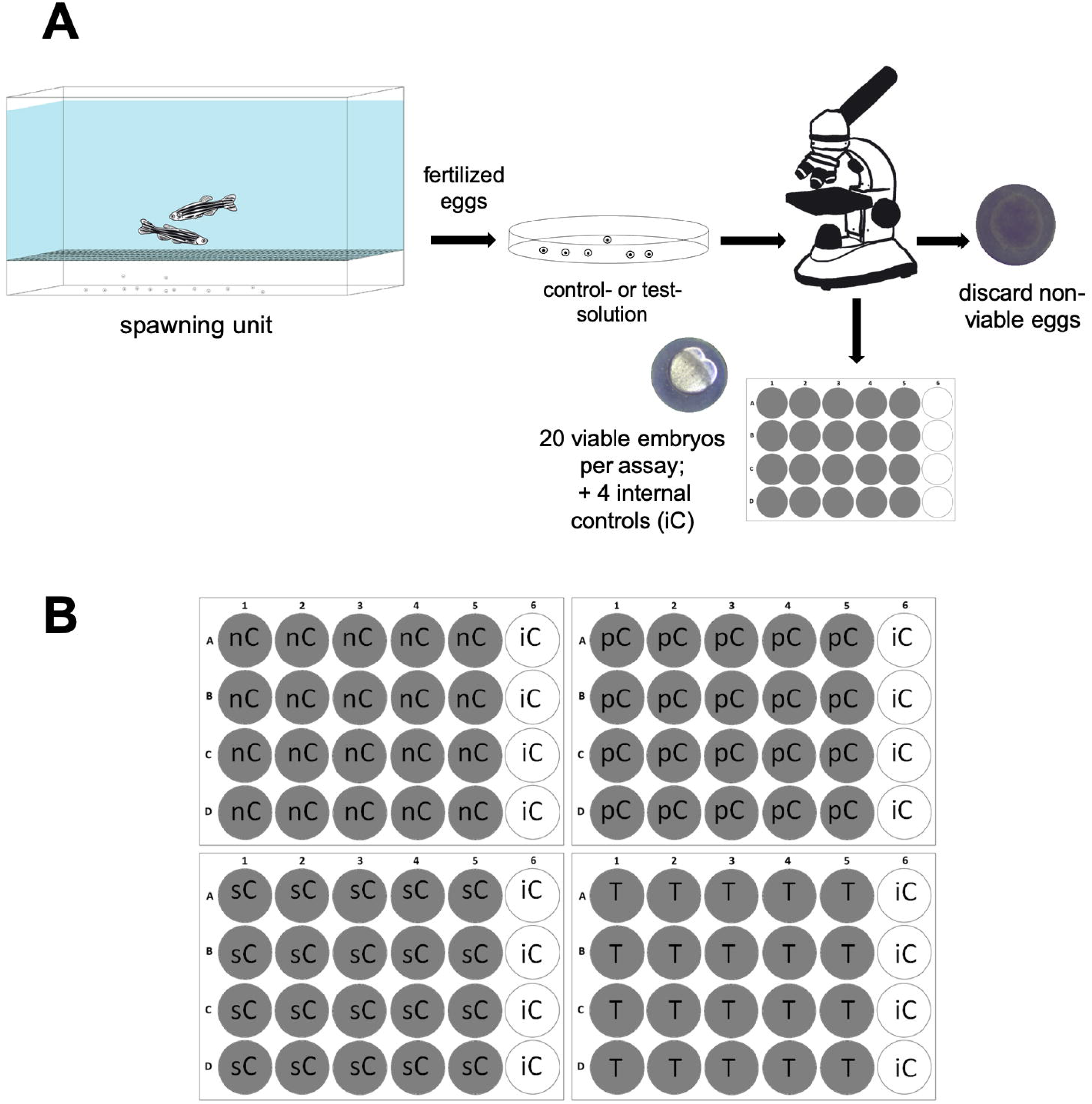
Setup of the zebrafish embryo development assay. A: Freshly fertilized eggs are transferred into a petri dish containing the test solution, viable eggs are selected microscopically, and 24 eggs containing viable embryos are transferred into a 24 well plates (1 egg per well). 20 wells contain 1 ml of BKI solution at a defined concentration (T), 4 wells serve as internal control (iC); **B:** 24 well-plate setup. T = BKI test concentration; nC = negative control (E3 medium/osmosis water); iC = internal control (E3 medium/osmosis water); sC = solvent control (DMSO); pC = positive control: dichloroaniline (4 g/L in water) (adapted from [31; 34])

**Figure 2.**
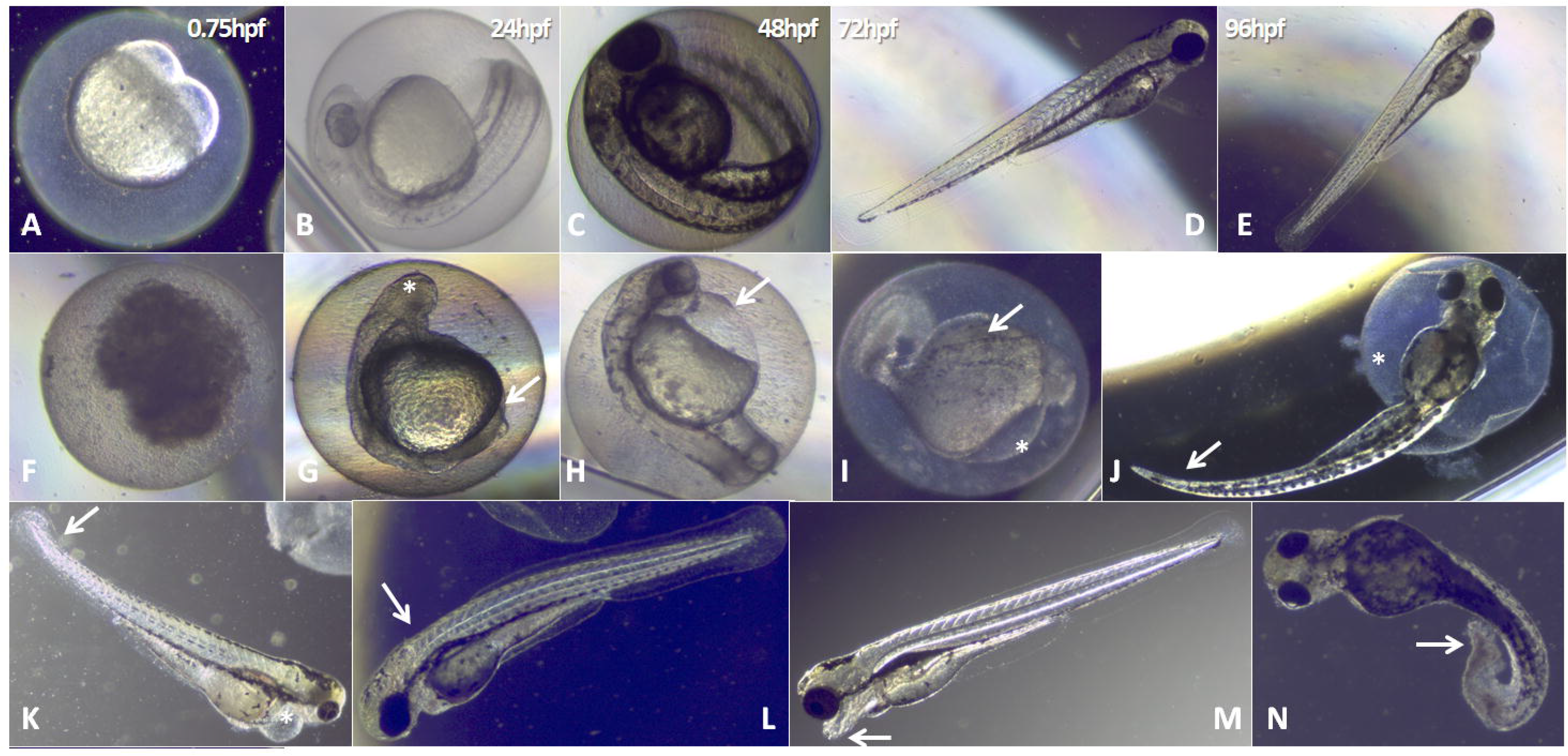
Normal development of the zebrafish embryo and BKI-induced malformations observed during test period. A-E: normal zebrafish embryo development at 0.75, 24, 48, 72 and 96h post fertilization. F-N show representative images of embryonal alterations that were observed during BKI-exposure. F: coagulated embryo (observed with BKI-1796, 24hpf/50μM); G: absence of eye bud (*) and tail non-detachment(→) (BKI-1796, 24h/50 μM); H: pericardial edema (→) (BKI-1517, 24h/50μM); M and I: body deformation(→)and pericardial edema(*) (BKI-1757, 24h/5 μM); J: tail curvature(→)and half-hatched embryo (*) (BKI 1517, 24h/20μM); K: curved tail (→) and pericardial edema (*) (BKI-1796, 24h/40μM); L: back curvature (scoliosis→) (BKI-1750, 24h/10 μM); M: head deformation(→) (1796, 24h/10μM); N: curved tail(→)(BKI-1796, 24h/10 μM).

### 2.7 Proposed algorithm for the assessment of drug impact on zebrafish embryo development

The formula for calculating the impact score (S_i_) of a given drug or control solution is provided in Fig. 3. A total of 20 eggs (N_e_ = 20) were microscopically assessed for each concentration of a given compound or the respective negative or solvent controls. In each assay, a score of −1 was assigned for death during the 96hpf, each malformation seen on a hatched embryo received a score of −0.5, including non-hatched embryos at 96hpf. Thus, the score for a given assay (S_assay_) was calculated by subtracting the scores for dead embryos (N_d_) and the scores for malformations (N_m_ divided by half) from N_e_ (see Fig. 3). The mean control score (S_mean_) was determined by calculating the mean of the negative control score (S_neg_) and the solvent control score (S_DMSO_). The overall impact score for a given drug concentration (S_i_) was finally calculated by subtracting S_mean_ from the S_assay_ achieved with the test dilution. We propose that a negative S_i_ indicates interference, and a S_i_ of 0 or higher implies no interference in early embryo development.

**Figure 3.**
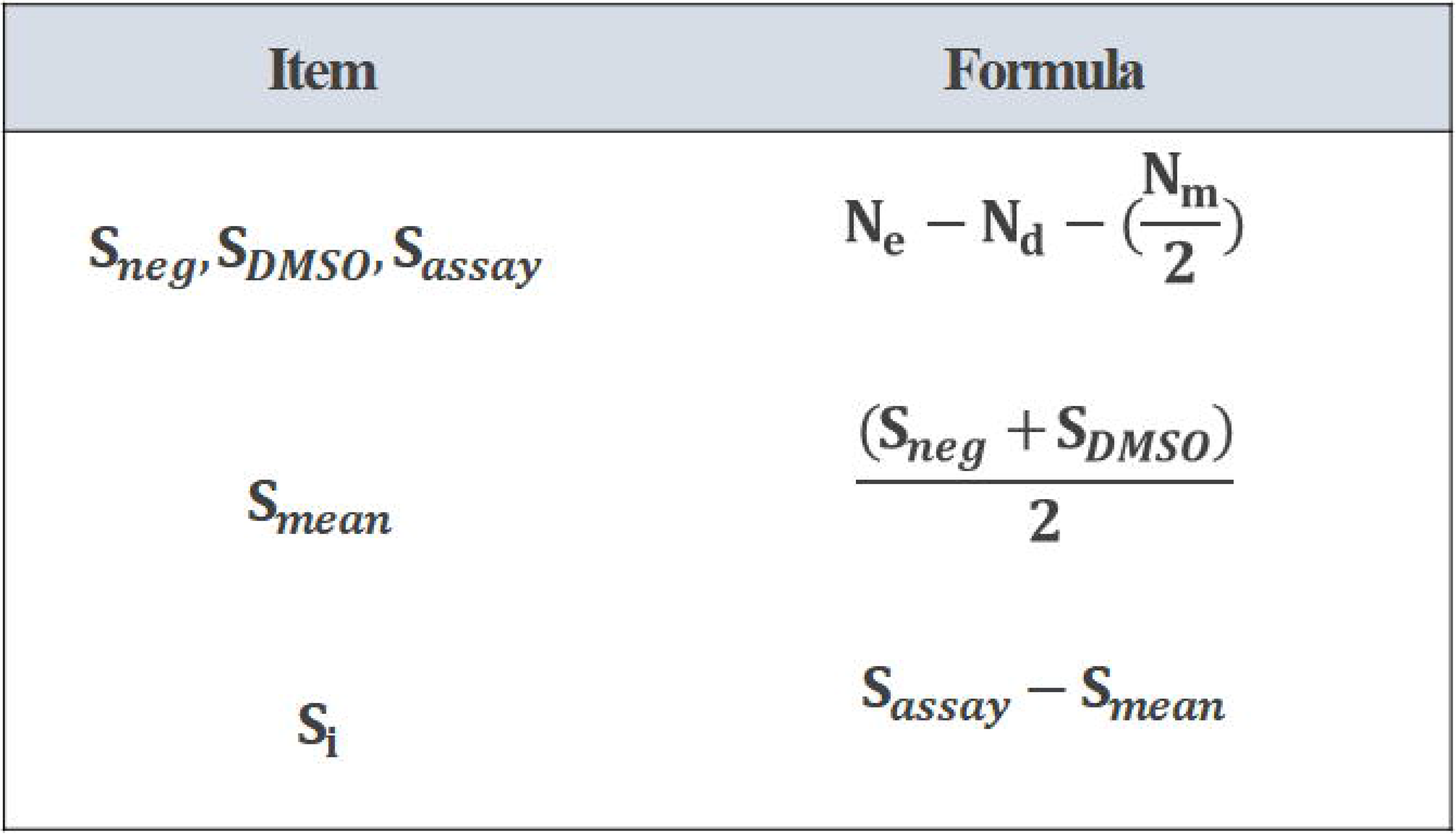
Proposed algorithm for the zebrafish embryo development assay score calculation. Every compound concentration receives a score based on the negative control and the solvent control. S_neg_ = negative control score (no compound, no DMSO); S_DMSO_ = solvent control score (no compound, 0.01% DMSO); S_assay_ = assay score (score for each compound concentration); S_mean_ = control mean score; S_i_ = impact score; N_e_ = number of embryos introduced per test item (20 per compound and concentration); N_d_ = number of dead embryos after 96hpf (score = −1 for each dead embryo); N_m_: number of embryonal malformations observed after 96hpf (score = − 0.5 for each malformation, including non-hatched embryos at 96hpf).

## RESULTS

### Effects of BKIs on Neospora caninum tachyzoites and mammalian cells cultured in vitro

The structures of the BKIs assessed in this work are depicted in Table 1. When added concomitantly to infection of HFF monolayers by *N. caninum*, they inhibited proliferation of tachyzoites with EC_50_s ranging between 0.035 and 0.180μM (Table 1 and supplementary Figure 1). Compounds were not toxic for HFF host cell monolayers at a concentration of up to 20μM, and no cytotoxicity was seen up to 40 or 80 μM in CRL-8155 and HEPG2 cells when assessed by Alamar Blue assay, except for BKI-1553, with which impaired the viability of CRL-8155 cells to 50% at 39.5μM (Table 1).

### Effects of BKIs on mouse pregnancy and offspring viability

The results on pregnancy interference of 15 BKIs in pregnant mice are shown in Table 2. In the placebo group, 4 of 6 mice undergoing mating became pregnant, with a total of 19 pups born alive, and no deaths observed during the follow-up period of 14 days. None of the 15 BKIs assessed in these experiments notably affected non-pregnant mice (data not shown). In pregnant mice, 5 compounds did not interfere in pregnancy outcome, while the others showed moderate to dramatic adverse effects in terms of offspring survival.

**Table 2.** Results of the mouse pregnancy interference tests and zebrafish embryo development interference tests using a panel of 15 BKIs. With the exception of BKI-1796, drugs were applied by daily oral gavage at a dose of 50 mg/kg/day during 5 days from days 7-11 of pregnancy. Fertilized zebrafish eggs were exposed to 6-8 different drug concentrations (0.2 – 50 μM) until 96h post fertilization. Grey background highlights those compounds for which the two models show corresponding results.

| Treatment | BALB/c mice |  |  | Zebrafish embryos |
| --- | --- | --- | --- | --- |
| | Fertility | Pups/dams | Offspring mortality | Lowest concentration exerting a negative $S_i$ (μM) |
| Placebo | 4/6 | 19/4 | 0/19 | n.a. |
| 1294 | 4/6 | 27/4 | 0/27 | 40 |
| 1369 | 5/6 | 31/5 | 0/31 | 40 |
| 1770 | 3/6 | 17/3 | 0/17 | >50 |
| 1750 | 4/6 | 19/4 | 0/19 | 5 |
| 1772** | 3/6 | 17/3 | 0/17 | 2 |
| 1553 | 3/6 | 16/3 | 6/16 | 5 |
| 1660 | 3/6 | 14/3 | 3/14 | 5 |
| 1713 | 4/6 | 20/4 | 4/20 | 20 |
| 1774** | 4/6 | 22/4 | 3/22 | 20 |
| 1757 | 4/6 | 15/4 | 3/15 | 0,2 |
| 1517 | *2/7 | 0/7 | -- | 0,2 |
| 1796<sup>a</sup> | *4/6 | 0/4 | -- | 0,2 |
| 1751 | *2/6 | 0/2 | -- | 5 |
| 1649 | 4/6 | 0/4 | -- | 10 |
| 1748 | 3/6 | 5/3 | 1/5 | 20 |

<sup>a</sup> BKI-1796 was applied at 25 mg/kg on days 1, 3 and 5
\* dams were determined to be pregnant by weight gain in the last trimester of pregnancy. They lost their offspring shortly before or during birth.
\*\* indicates compounds where precipitation was noted visually in water at concentrations of 20 μM and above.

Treatment with BKI-1294 resulted in 4 of 6 pregnant mice giving birth to an overall number of 27 pups, of which all survived until 14 days post-partum. Similar results were obtained with BKI-1369. Relative to the placebo control, BKI-1750 resulted in the exact same fertility and number of pups. BKI-1770 and BKI-1772 were only slightly lower for fertility and this difference is not significant in comparison to the controls.

In the cases where interference in pregnancy was noted, pups were either resorbed, aborted, born dead, or neonatal mortality took place within 2 days after birth. No further deaths occurred during the 12 days follow-up. Pup mortality rates between 15-37% were noted in experimental groups treated with BKI-1553, BKI-1660, BKI-1713, BKI-1757 and BKI-1774.

Other compounds interfered in pregnancy much more severely, such as BKI-1517. Abortion or a high degree of offspring mortality was also noted for BKI-1748. In the group treated with BKI-1751, 2 of 6 were supposedly pregnant as assessed by weight, but no pups were found on the day of supposed birth, and one pregnant mouse was found dead on day 21 post-mating. In the group treated with BKI-1796, which was assessed at 25mg/kg on days 1, 3 and 5, 4 out of 6 mice were supposed pregnant according to their weight gain, but no viable pups were born and blood stains were found in the cage. The fact that these animals lost up to 5g within 2 days indicates they had aborted.

### Effects of BKIs on early zebrafish embryo development

The assessment of effects of BKIs on zebrafish embryo development is based on the daily microscopic inspection of the 20 individual fertilized eggs for each drug and respective concentrations cultured 24-96hpf, either in the presence or absence of compounds. This visual assessment was blinded with respect to treatment. The typical morphological features of normally developing zebrafish embryos within those 96hpf are shown in Fig. 2A-E. No differences could be detected between the negative control group and DMSO control group. Typical examples of the different types of drug-induced malformations rated in our studies are shown in Fig 2.F-N.

In cases where toxicity was observed, the effects were mostly dose-dependent. The lowest concentrations causing adverse effects in early zebrafish embryo development for each compound are shown in Table 2. A detailed account of the number of malformations, deaths and impact scores obtained in the test groups are provided in supplementary Table 1, and for selected compounds, the results are also shown as graphs, with S_i_s plotted against drug concentrations as supplementary Figure 2. Additionally, supplementary Figure 3 shows graphs of the toxicity of selected BKIs versus time (hpf). Notably, only three compounds, namely BKI-1294, BKI-1369 and BKI-1770 exhibited very low embryo toxicity, with negative impact scores assigned to them at 40 and/or above 50μM. These three BKIs also do not interfere in pregnancy outcome in the mouse model (see above). Nine BKIs interfered in early embryo development at concentrations ranging from 2 to 20μM. Three BKIs (BKI-1757, −1517 and −1796) were very embryo-toxic at the lowest concentration of 0.2μM. BKI-1757 partially interfered in pregnancy outcome (3 neonatal deaths of 15 pups), while the other two exhibited more dramatic effects, with no viable offspring born.

## Discussion

The BKIs assessed in this study all exhibited promising *in vitro* activity against *N. caninum* tachyzoites, with IC_50_ values in the lower sub-micromolar range, with no viability impairment of tested mammalian cells. The lack of host cell impairment by these BKIs is an important prerequisite to advance studies in small animal models. Based on their *in vitro* efficacy and specificity, these drugs were promising leads as *N. caninum* therapeutics and for the treatment of other diseases caused by apicomplexan parasites.

Vertical transmission is the most important mode of *N. caninum* infection. Thus, compounds to be applied against neosporosis could be used in the treatment of pregnant animals when immunity is down-modulated to prevent exogenous transplacental transmission, and to limit endogenous transplacental transmission in chronically infected animals due to recrudescence. Compounds could also be applied to treat congenitally infected newborns to prevent the formation of tissue cysts [3]. Thus, besides reaching therapeutic levels in the CNS, drugs should be able to cross the placenta in order to eliminate infection in fetal tissues, without inducing harmful maternal physiological changes or exerting teratogenic effects, which most likely occur during the initial phases of embryo development due to rapid cell proliferation and increased metabolic activity [35].

While neosporosis is a disease that is mainly important in ruminants, economic considerations demand that studies on the basic features of *N. caninum* infection, potential vaccine candidates, and novel anti-parasitic drugs, to be carried out in small laboratory animals such as mice. Regardless of the drawbacks of using mouse models, even though they do not fully reflect *N. caninum* infections in cattle, they do provide initial information on protective effects of potential treatments [2].

The 15 BKIs assessed in this study can be divided into three distinct groups. Group 1 contains 5 BKIs that did not induce adverse effects in pregnant mice. Three of those (BKI-1294, 1369 and 1770) affected zebrafish embryo development only at high (40 μM or even higher concentrations), while the other two (BKI-1750 and 1772) exhibited zebrafish embryo-toxicity already at 5 and 2 μM, respectively. Group 2 is comprised of 5 BKIs that exhibit moderate interference in pregnancy with 15-37% of offspring loss within the first few days after birth. Zebrafish embryo development was impaired at 0.2μM by BKI-1557, and at 5 or 20μM by the others. Group 3 consisted of 5 BKIs that displayed fetal loss during pregnancy, abortion, still birth and neonatal mortality, with very few offspring still alive after birth. Zebrafish embryo development was impaired already at 0.2μM by two of these BKIs; the others exhibited effects on zebrafish embryos at concentrations between 5 and 20μM.

This limited data set illustrates two points: (i) the three compounds that exhibited interference in zebrafish embryo development at 0.2μM also had a negative impact on pregnancy outcome as reflected by the reduced offspring survival, indicating that embryo toxicity at 0.2μM is predictive for pregnancy failure in this mouse model; (ii) the lack of microscopically visible embryo toxicity does not imply that there will be a successful pregnancy outcome, suggesting that other determinants of maternal physiology, possibly affected by drug treatments, are important. However, it is also possible that visual inspection of morphological development cannot fully explain all mechanisms of embryotoxicity. Thus, more studies need to be carried out to corroborate these findings.

BKIs-1294, −1369 and −1770 are group 1-compounds not affecting zebrafish development at high concentrations. BKI-1294 was shown previously to be safe and efficacious in non-pregnant and pregnant BALB/c mice infected with *N. caninum* tachyzoites [22,25], was also highly efficacious in non-pregnant *T. gondii* infected mice [19], and in pregnant mice infected with *T. gondii* oocysts [20]. However, the latter model was based on CD1 outbred mice, and *T. gondii* infected and drug treated CD1 mice gave birth to significantly lower numbers of pups compared to non-treated CD1 mice. This implied that interference in pregnancy could be an issue dependent on the mouse strain, in addition to other factors. Recently BKI-1294 treatment was applied in pregnant sheep experimentally infected with *T. gondii* oocysts where it was partially efficacious against *T. gondii* infection, and no adverse effects of drug treatment were noted in this small ruminant model [36]. BKI-1369 and BKI-1770 are lead compounds for the treatment of diarrhea caused by cryptosporidiosis in calves [15,17]. In addition, excellent BKI-1369 efficacy was demonstrated against *Cystisospora suis* infection in piglets [18], and in a pig model of acute diarrhea caused by *Cryptosporidium hominis* ([16].

BKI-1553, a compound with low hERG inhibitory activity [24], also exhibited considerable effects on pregnancy outcome when applied at 50 mg/kg, but it had been shown earlier that this effect could be alleviated by a three times treatment at 20 mg/kg every second day, which reduced neonatal mortality but still maintained considerable anti-parasitic activity in *N. caninum* infected pregnant mice [23]. BKI-1553 is highly active against *T. gondii in vitro*, crosses the blood-brain barrier in mice when orally applied, and reduces the *T. gondii* burden in brain, lungs, and liver [24]. The safety and efficacy of BKI-1553 was also demonstrated in pregnant sheep experimentally infected with *N. caninum* tachyzoites. Subcutaneous application resulted in reduced fetal mortality by 50% [21]. *N. caninum* was abundantly found in placental tissues, but parasite detection in fetal brain tissue decreased from 94% in the infected/untreated group to 69-71% in the treated groups.

Three BKIs exhibited clear embryo-toxicity at the lowest dose of 0.2μM, and also induced moderate to severe disturbances in pregnancy outcome when applied at 50mg/kg for 5 days. Among them, BKI-1517 has been previously investigated in *Neospora*-infected pregnant mice. At 50mg/kg, 100% neonatal mortality was observed, and this was confirmed in this study. At 20mg/kg, BKI-1517 significantly decreased the rate of vertical transmission and increased offspring survival [23]. BKI-1517 also displayed promising efficacy against *Cryptosporidium* spp. [12, 14]. In our experiments, the compound did not show toxicity against mammalian cells. However, it exhibited high toxicity at all concentrations in the zebrafish assay, which explains its detrimental effects in pregnant mice.

The reasons for the toxicity of some of the BKIs is not known. BKIs act as ATP competitive inhibitors of CDPK1 [37], but interactions with other kinases exhibiting bulkier gatekeeper residues cannot be ruled out. One of the possible off-targets is PKD3, another serine/threonine kinase, which is associated with the mitotic apparatus, including spindle, centrosome, mid-body formation and development of cardiac and skeletal muscle tissue [38, 39]. The dual-specificity kinase MEK1 is another kinase potentially targeted by BKIs. It plays an activating role in the ERK/MAPK cascade, which is involved in cell-fate determination and regulation of cell proliferation and differentiation. MEK1 gene disruption in embryonic stem cells resulted in death of embryos after 10 days of gestation [40]. These results suggest that both kinases are important for the developing fetus, and compounds that inhibit their normal function may result in teratogenic effects.

While the mouse pregnancy safety model plays an important role in drug development, especially for BKIs targeting vertical transmission of pathogens, it cannot be maintained as a routine screening platform due to the extensive use of animals, the high amount of compound required, time-consuming procedures, and the costs involved. In addition, there are physiological differences between mouse placenta and the placenta of humans and other mammalian species, with anticipated consequences regarding fetal drug susceptibility issues [41]. In this respect, the zebrafish embryo development model could serve an important purpose for the identification of BKIs with teratogenic potential, with higher throughput, lower costs and more rapid results, and could potentially serve to eliminate embryo-toxic compounds early on. One important limitation is that feasibility is restricted to water-soluble compounds. We observed precipitates in tests with BKI-1772 and BKI-1774 at 20 μM and above, and respective results at 20 μM are inconclusive. In addition, it only makes sense to carry out such studies at therapeutic concentrations. Thus, it is helpful to know the pharmacokinetics of each compound, and then choose the concentrations accordingly. Despite these potential pitfalls, the zebrafish model can serve as an important pre-screening tool that could contribute to the 3Rs (replacement, reduction, and refinement) and allow more efficient testing of BKIs, as well as other anti-parasitic drugs to be applied during pregnancy. In addition, future studies with a much larger dataset will show, how our proposed algorithm for the calculation of a drug impact score S_i_ in zebrafish embryo development will perform in predicting drug interference in pregnancy in mice.

## ACKNOWLEDGEMENTS

We are grateful to Vreni Balmer, Adriana Aguado-Martinez, and Naja Eberhard for their help and valuable input and help in parasite cultivation and zebrafish embryo maintenance.

## DECLARATIONS

**Funding:** This study was financed by the Swiss National Science Foundation (SNSF) grant 310030_184662, and the National Institutes of Health (NIH) grants R01AI089441, R01AI111341, R21AI123690, and R21AI140881.

## Ethical Approval

All protocols involving animals were approved by the Animal Welfare Committee of the Canton of Bern under the license BE101/17.

