## Supplemental Figure 1 for "Comparative assessment of the effects of bumped kinase inhibitors on early zebrafish embryo development and pregnancy in mice"

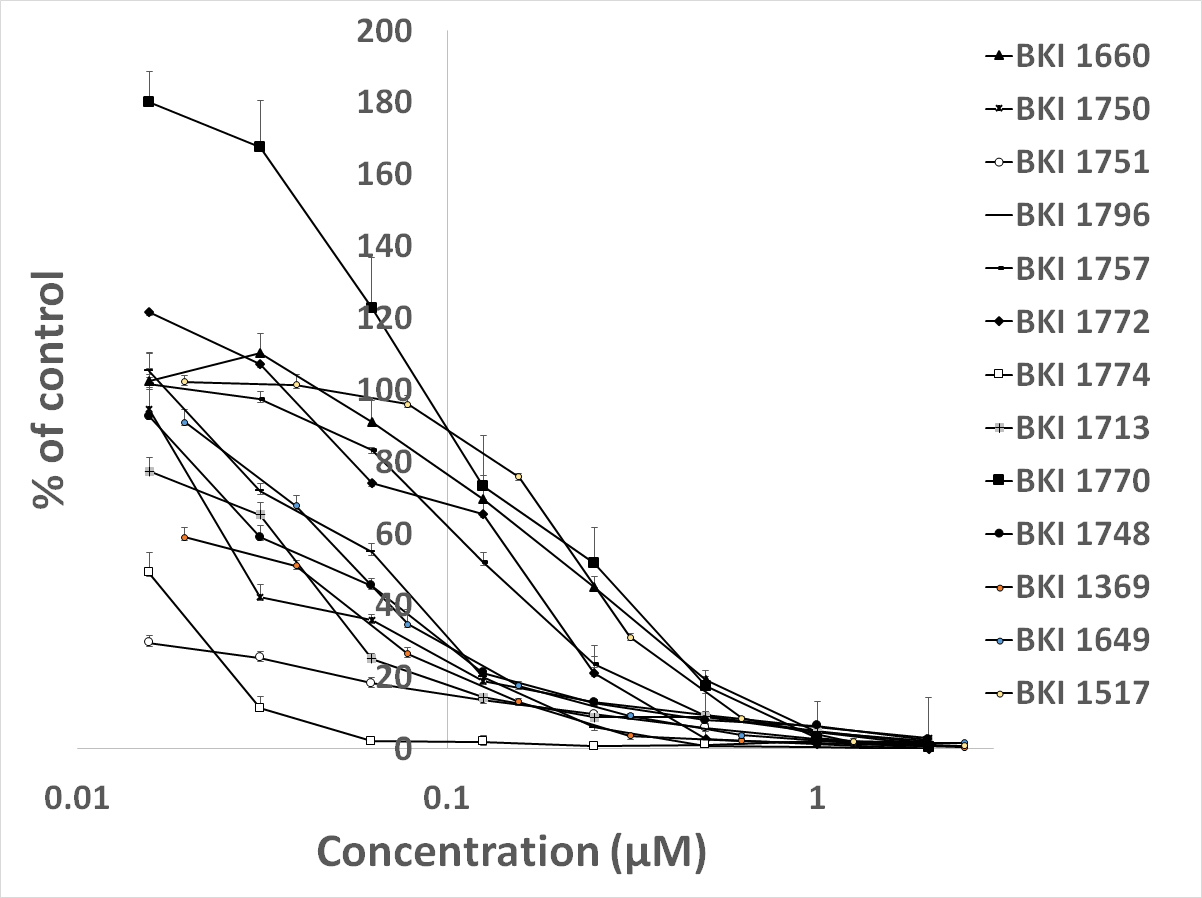

**Supplementary Figure 1:** Dose-response curves for 13 BKIs, demonstrating dose-dependent inhibition of *N. caninum* tachyzoite proliferation. Data on BKI-1294 and BKI-1553 has been published elsewhere ([25] and [23] respectively).
