## Supplemental Figure 2 for "Comparative assessment of the effects of bumped kinase inhibitors on early zebrafish embryo development and pregnancy in mice"

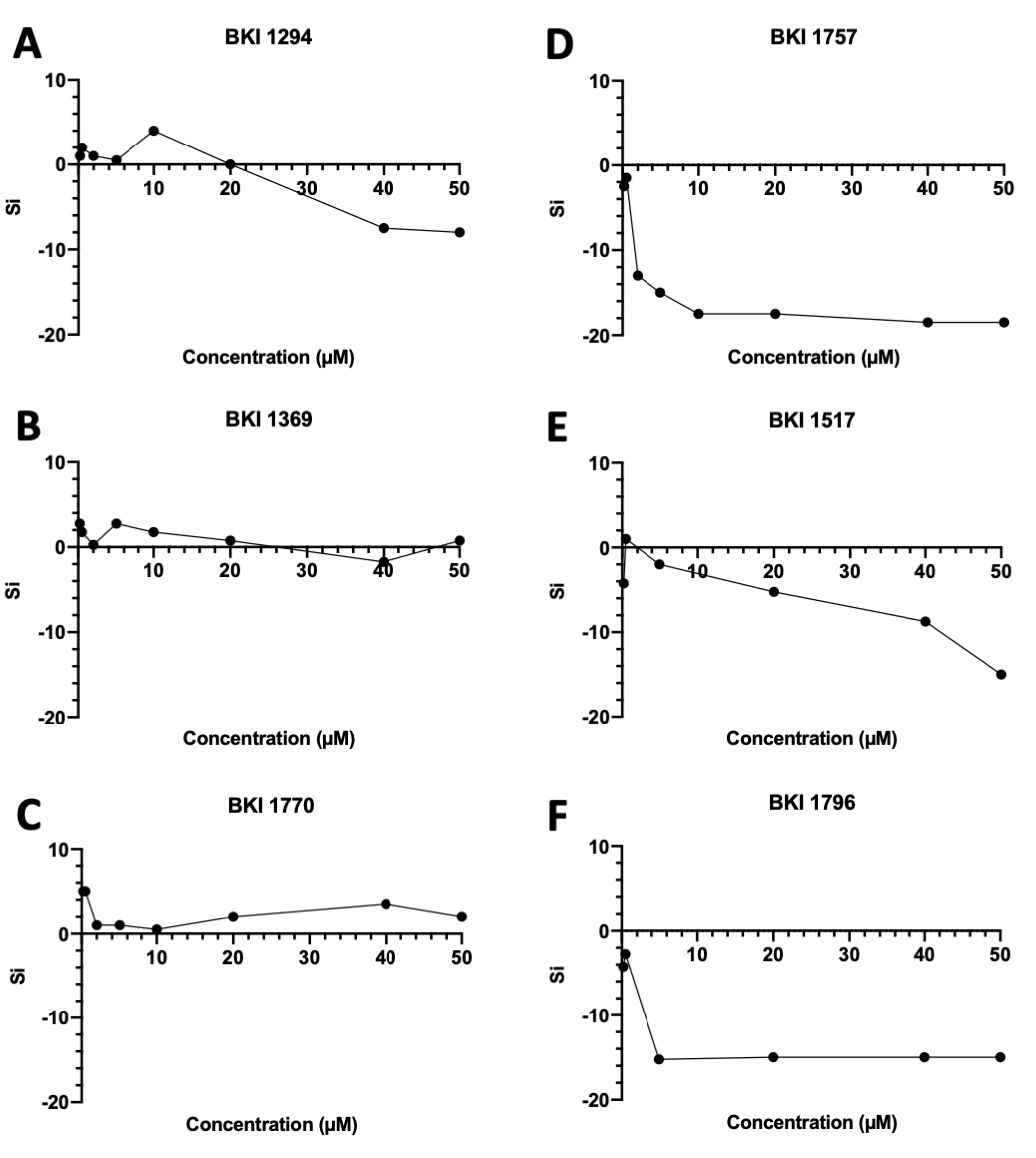

**Supplementary Figure 2:** Graphs depicting the impact of selected BKIs on zebrafish embryo development. A = BKI-1294; B = BKI-1369; C = BKI-1770; D = BKI-1757; E = BKI-1517; F = BKI-1796. For each compound the calculated impact Score S_i_ is plotted against the drug concentration.
