## Supplemental Figure 3 for "Comparative assessment of the effects of bumped kinase inhibitors on early zebrafish embryo development and pregnancy in mice"

**Supplementary Figure 3:** Graphs depicting the impact of selected BKIs on embryo survival. A = BKI-1294; B = BKI-1369; C = BKI-1770; D = BKI-1757; E = BKI-1517; F = BKI-1796. For each drug and drug concentration the numbers of still living embryos on the y-axis are plotted against hours post fertilization (0, 24, 48, 72 and 96 hpf). Negative controls and solvent controls (0.01% DMSO) are also shown for each experiment. Malformations are not included.
